# *De novo* long-read assembly of a complex animal genome

**DOI:** 10.1101/187054

**Authors:** David Eccles, Jodie Chandler, Mali Camberis, Benard Henrissat, Sergey Koren, Graham Le Gros, Jonathan J. Ewbank

## Abstract

Eukaryotic genome assembly remains a challenge in part because of the prevalence of complex DNA repeats. This is a particularly acute problem for holocentric nematodes because of the large number of satellite DNA sequences found throughout their genomes. These have been recalcitrant to most genome sequencing methods. At the same time, many nematodes are parasites and some represent a serious threat to human health. There is a pressing need for better molecular characterization of animal and plant parasitic nematodes. The advent of long-read DNA sequencing methods offers the promise of resolving complex genomes. Using *Nippostrongylus brasiliensis* as a test case, applying improved base-calling algorithms and assembly methods, we demonstrate the feasibility of *de novo* genome assembly matching current community standards using only MinION long reads. In doing so, we uncovered an unexpected diversity of very long and complex DNA repeat sequences, including massive tandem repeats of tRNA genes. The method has the added advantage of preserving haplotypic variants and so has the potential to be used in population analyses.

## Introduction

Human hookworm infections by the parasitic nematodes *Necator americanus* and *Ancylostoma duodenale* continue to be a major global health problem. Next generation sequencing techniques open the door to molecular epidemiological monitoring of nematode and helminth parasites in endemic areas. Such studies are, however, hampered by the heterogeneous nature of parasite populations and by the intrinsically complex genome structures of nematodes [1]. The highly portable nature of MinION sequencers makes them well-suited for field studies, and their capacity to generate long sequence reads offer the promise of overcoming both these obstacles. *Nippostrongylus brasiliensis* is a gastrointestinal nematode that infects rodents. It is widely used as a model for human hookworm (e.g. ref. [2]). Previous attempts to assemble the *N. brasiliensis* genome using short DNA reads have resulted in a highly fragmented sequence (**Table 1**); almost 30% of predicted protein coding genes (6276/22796) are on contigs that are less than 10kb long. We took *N. brasiliensis* as a test case to evaluate the possibility of generating a genome sequence *de novo* from a heterogeneous population.

**Table 1.** Genome and transcriptome contiguity scores. Statistics for the current WTSI reference sequence are given for comparison.

| Description | Size (kb)<br>[Contigs] | N50 (kb)<br>[L50] | N90 (kb)<br>[L90] | Min (kb) | Max (kb) |
| --- | --- | --- | --- | --- | --- |
| WTSI reference | 294400<br>[29375] | 33.5<br>[2024] | 4.3<br>[11638] | 0.5 | 394.2 |
| WGA, Canu only | 119196<br>[3280] | 53.1<br>[647] | 17.3<br>[2199] | 1.8 | 388.2 |
| Canu only* | 347186<br>[3583] | 209.2<br>[415] | 38.9<br>[1964] | 1.7 | 2048.3 |
| Trinity [*de-novo*] | 180448<br>[291671] | 0.8<br>[53613] | 0.3<br>[219557] | 0.2 | 21.5 |
| Trinity [genome-guided] | 249065<br>[352994] | 1.2<br>[50569] | 0.3<br>[245765] | 0.2 | 22.0 |
| Trinity [Expression-filtered] | 71457<br>[52302] | 2.0<br>[11072] | 0.7<br>[34924] | 0.2 | 22.0 |

\* Local read correction has minimal effect on the contiguity statistics

*de novo* genome assembly based on long DNA reads relies on either hybrid strategies incorporating, for example, short-read DNA sequencing [3-6] or non-hybrid long-read only methods [7-11] (see [12] for a review). There have now been successful chromosome-scale assemblies of large genomes, primarily using the PacBio single-molecule realtime (SMRT) sequencing platform to generate contigs (e.g. [13]), combined with long-range linking information (e.g. [14, 15]) but the approach can still be challenging. We therefore chose a simpler system to evaluate the feasibility of assembling a genome for *N. brasiliensis* using improved analysis methods.

## Results and Discussion

Whole-genome amplification was used to generate DNA from a single adult worm. This was then sequenced with a single run on a MinION Mk1b (see Methods for details). The original base-calling produced 1722835 reads totaling 4.9 Gb of sequence. These reads were processed using a default Canu [16] v1.4 assembly (including error correction and read trimming) giving an unpolished assembly of 4581 contigs for 117 Mb of assembled genomic sequence (**Table 1**). We then re-called the same raw nanopore data using Albacore, the production base-caller that recently implemented a transducer algorithm for homopolymer detection, previously a persistent limitation to analysis of MinION reads [17]. Overall, we obtained 2074871 called reads for 6.5 Gb of sequence. This represents a substantial increase ( >30%) in length over the previous version. These reads were fed into an up-dated version of Canu (v1.5), with improved read correction and consensus calling, and more accurate graph information in the assembly output, giving an unpolished assembly of 3280 contigs and 119 Mb of assembled genomic sequence. Not only were there local improvements in the calling of homopolymer sequences (**Figure 1A**), but the contigs showed a much improved length distribution, and a much higher proportion of graphs were resolved (**Figure 1B, C**), in part because of a improved resolution of repeat sequences (**Figure 2A-C**).

**Figure 1.**
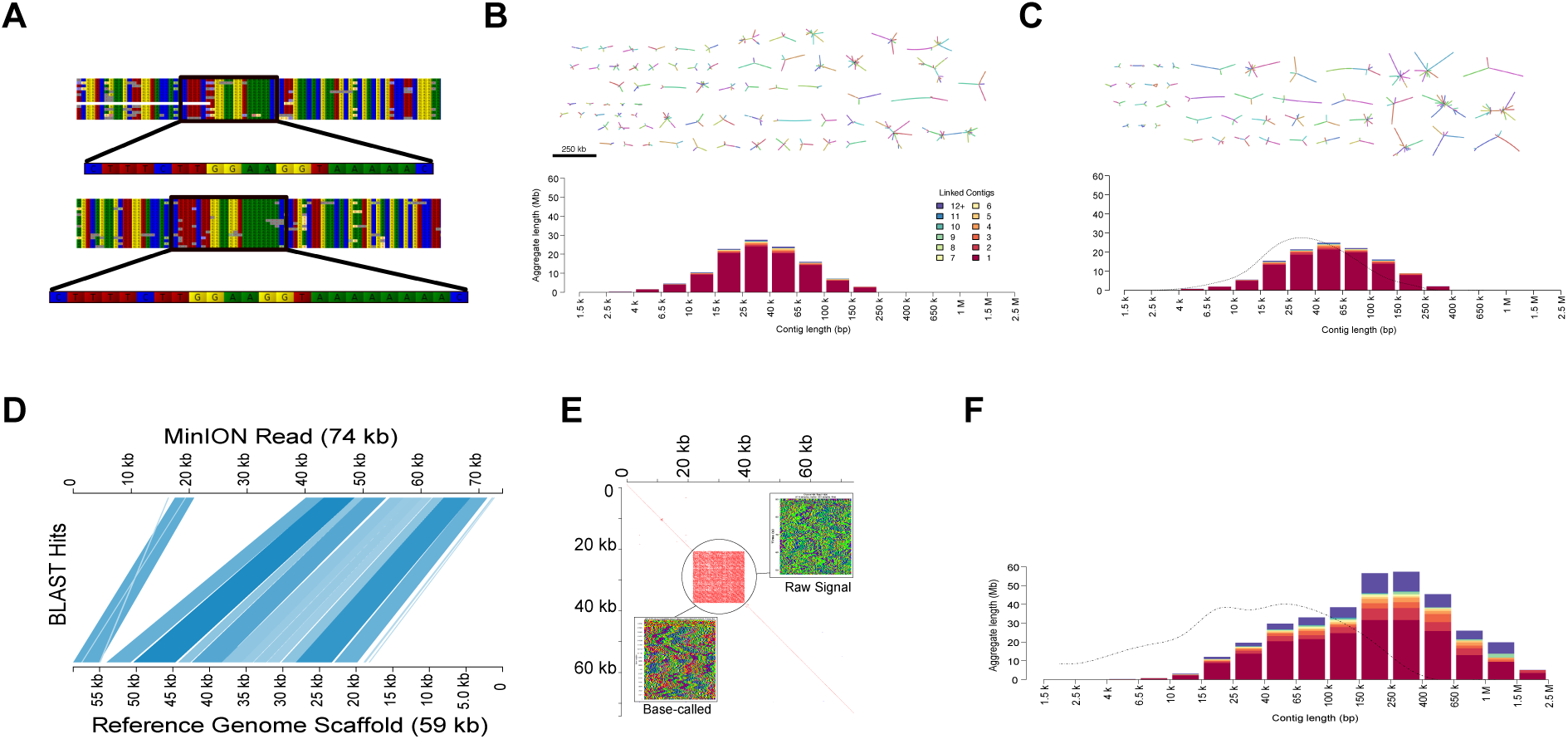
Improved sequence fidelity and assembly using updated methods. (A)Homopolymeric regions that were compressed in the original sequence (top) were expanded when the basecaller included a “transducer” mode that incorporates signal length into the called sequence (bottom). The consensus sequences for the boxed region of the aligned individual uncorrected reads are shown. Sanger sequencing of PCR amplicons confirmed the accuracy of the newer base‐calling. **(B, C)** Distribution of fragment sizes and complexity for contigs assembled using old **(B)** and new **(C)** methods with amplified DNA from a single worm. Graphs (drawn to scale as indicated in **(B)**), are also shown; the non‐branched graphs representing more than 90% of the sequence (dark magenta bars) have been omitted. The distribution of fragment sizes from **(B)** is shown as a dashed line in **(C)** for direct comparison. **(D)** Alignment of a single 74 kb read against the corresponding scaffold of the current WTSI reference genome, plotted using Kablammo [23]. **(E)** Identification of a very long stretch of complex tandem repeats (21 kb with a 171 bp repeat unit) within the same read. **(F)** Distribution of fragment sizes and complexity for the final assembly, compared with that of the WTSI reference genome (dashed and dotted line).

**Figure 2.**
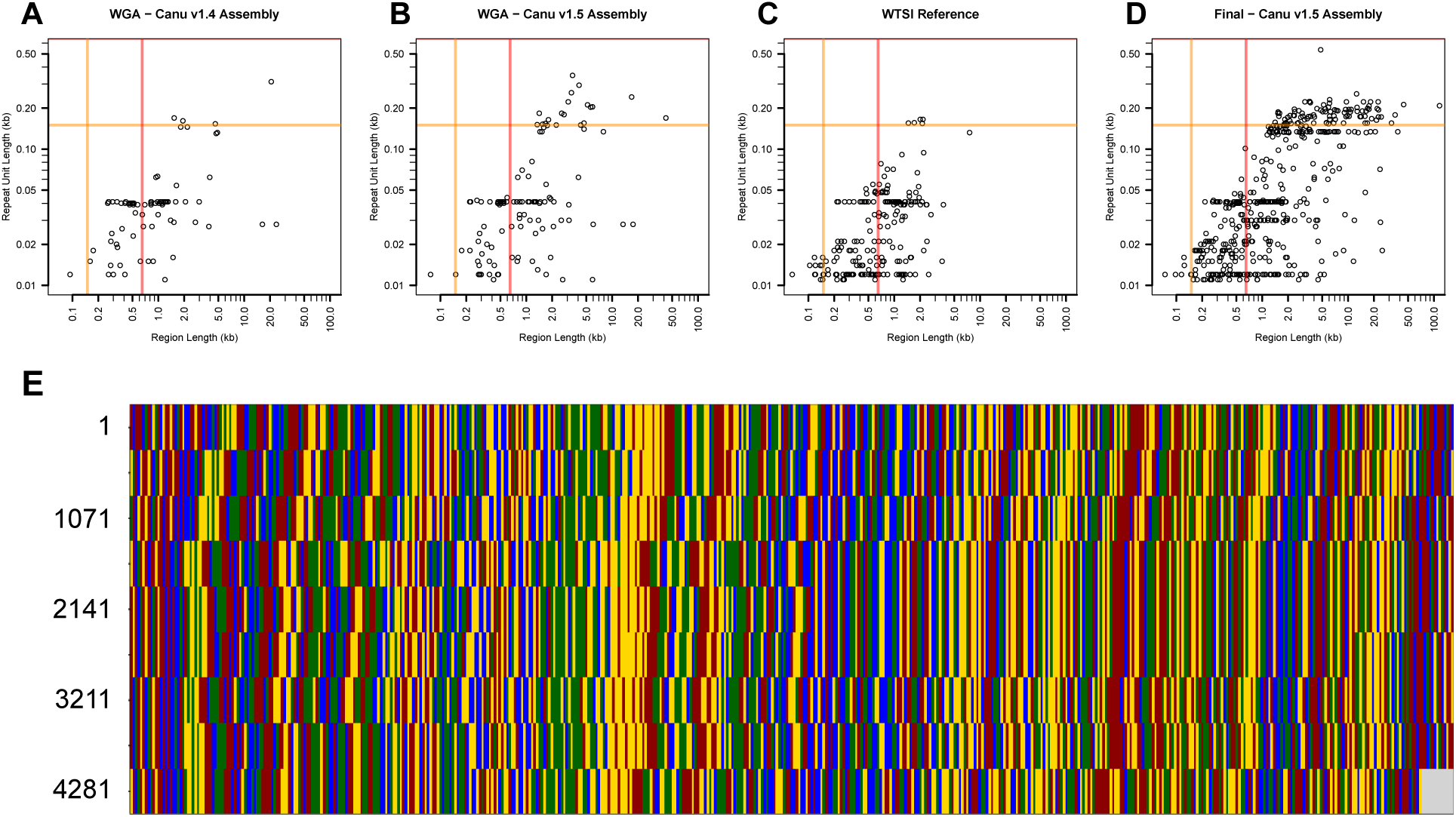
Analysis of repeat sequences within different assemblies. SATFIND [18] was used on contigs > 2.5kb. The total length of each region of repeated DNA sequence is plotted against the repeat’s unit length, for the assemblies from DNA amplified from a single worm (WGA), using Canu v1.4 (**A**) or Canu v1.5 (**B**), for the WTSI genome assembly (**C**) and our final assembly (**D**). The orange and red lines are at 150 bp (typical maximum read length for an Illumina HiSeq run) and 650 bp, respectively. Any VeCTRs with a unit length longer than 150 bp would not be identifiable as repetitive sequence on an Illumina sequencer. Any VeCTRs with a region length longer than 650 bp will be collapsed into a shorter region if only non‐mate‐paired 150 bp reads are used for the assembly. (**E**) Alignment of the region corresponding to the longest repeat unit length in (**D**), with each base represented as a colored line. Although the resolution of repeats is greatly improved compared to the WTSI assembly, such sequences still present a challenge for assembly, as evidenced by the fact that the 5’ end of this sequence corresponds to a contig end (tig00023164; coordinates on the left). The color code is as in Figure 1A.

These results encouraged us to sequence DNA from a population of worms resident in the small intestines of mice. We used 4 sample preparation methods that gave very different results both in terms of yield, which varied more than 2-fold in total, up to 4.7 Gb for a single run, and in read length distribution (**Figure 3**). In all cases, Albacore out-performed the older base-caller, although interestingly the improvement was far from uniform, varying from less than 10% to greater than 30% in total reads called (**Table 2**). As a further gauge of the accuracy of read calling, we compared a random subset of the longer reads with the current *N. brasiliensis* reference genome. While overall there was generally a good equivalence, especially given the potential genetic difference between the two samples (see Methods), there were numerous instances where the reads were substantially longer than the corresponding sequence in the reference genome. Upon further examination, some of these were revealed to reflect the presence of very long stretches of complex tandem repeats (or VeCTRs) that had been compacted in the WTSI reference genome (**Figure 1D, E**).

**Figure 3.**
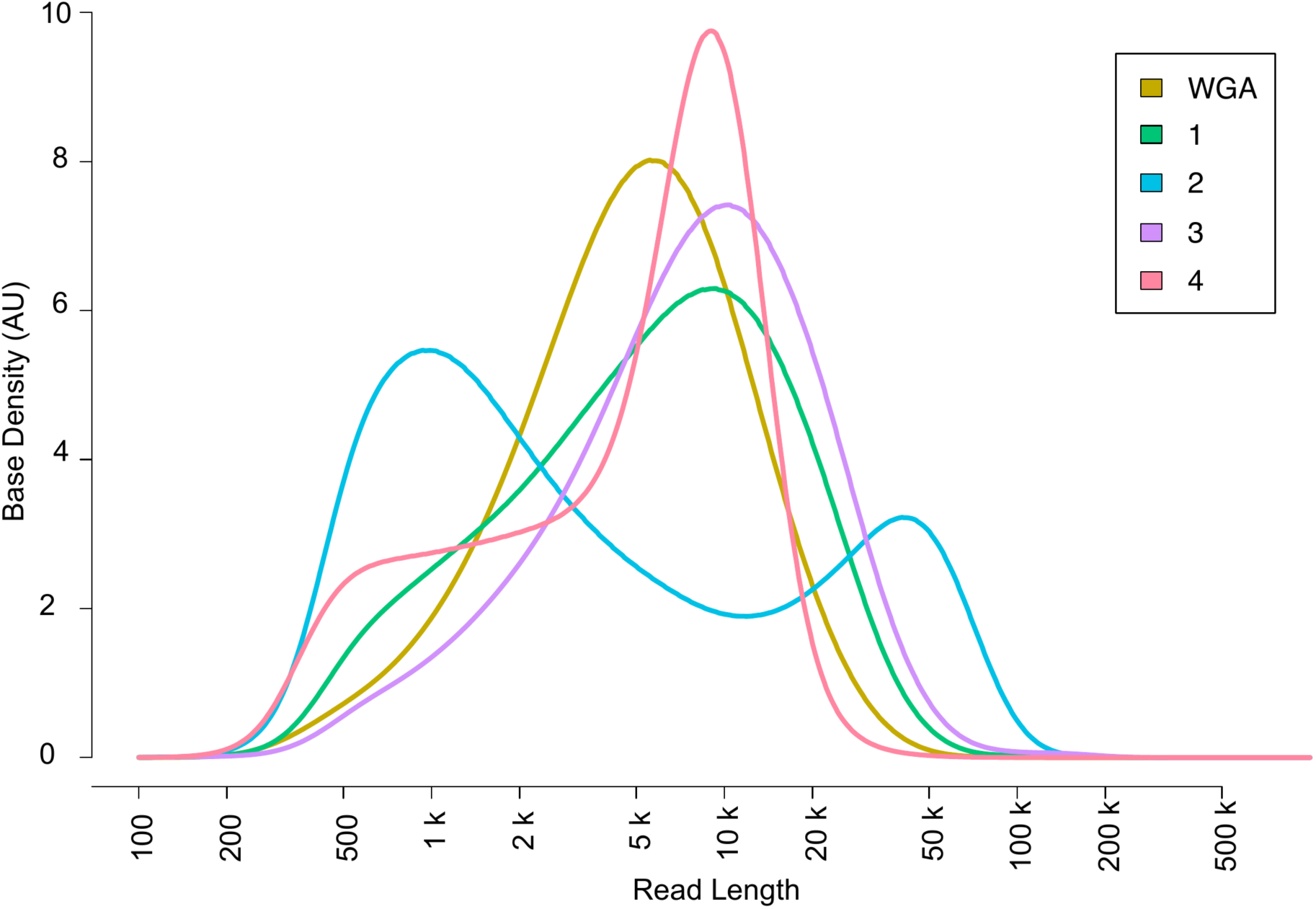
Effect of extraction method on the size of DNA reads. The distribution of read lengths for unamplified DNA extracted by each one of 4 methods (see Table S2). The curve for the sequences from the DNA amplified from a single worm (WGA) is shown for comparison.

**Table 2.** Yield of sequence using different extraction and analysis methods. The percentage of the raw reads that were included in the output of the 2 methods (MinKNOW and Canu 1.4, or Albacore and Canu 1.5) are shown together with the total sequence lengths. See Methods for details.

|  | Yield |  |  |  |  |
| --- | --- | --- | --- | --- | --- |
| Preparation Method | Raw reads | MinKNOW |  | Albacore |  |
|  |  | % | Gb | % | Gb |
| Single worm, WGA | 2093867 | 82.3 | 4.889 | 99.1 | 6.472 |
| 1. Sonication, DNeasy | 930957 | 99.9 | 2.375 | 99.1 | 2.603 |
| 2. Sonication, G2, Tip20 | 1845520 | 85.2 | 2.289 | 98.0 | 3.002 |
| 3. Direct lysis, DNeasy | 451578 | 99.9 | 1.723 | 98.8 | 2.028 |
| 4. Pipette squashed, G2, Tip20 | 2333064 | 83.9 | 3.834 | 98.4 | 4.678 |

We combined the reads from the 4 samples and fed them into Canu v1.5. This gave an assembly of close to 350 Mb, with a maximum contig length of >2 Mb and an N50 of 210 kb. A total of 251.6 Mb (72.5%) of the sequence, including all of the longest ( >1.5 Mb) contigs, was contained within 2594 non-branched sub-graphs of the Canu assembly graph, with 2413 of them made from a single contig, and the others containing an average of 2.15 linked contigs. The remaining 95.6 Mb was captured by a total of 780 contigs in 89 non-trivial sub-graphs. A small number of these sub-graphs had very complex structures (**Figure S1**). Inspection showed this to result from the presence of long non-tandem repeat sequences. Indeed this, together with real haplotypic sequence diversity (see below), was the most common cause of assembly ambiguity. Resolving such structures would require a greater depth of coverage and/or even longer reads.

The final assembly contained a rich diversity of repeat sequences (**Figure 2**). The repeat with the longest unit length (535 bp) corresponds to a region with 10 tandem copies (**Figure 2D**) of an 5S rRNA gene interspersed with an snRNA gene, the source of the spliced leader RNA that is added to many transcripts. The gene sequences and this arrangement are well-conserved in *Caenorhabditis elegans*, where these pairs are repeated over a region of 16 kb (V:17,118,000..17,132,000; WS260). In the WTSI assembly, on the other hand, equivalent sequences, in single copy, were found at contig ends. There were also transposon-associated repeats, the repeats that correspond to reiterated amino acid motifs in proteins, and the dispersed satellite DNA sequences found in holocentric nematodes [18]. With regard to the other VeCTRs, their length and the constituent repeated sequences are diverse. There was, for example, no sequence similarity among the 5 most compressible VeCTRs (**Figure S2**). The longest of them comprised close to 150 copies of a ca. 200 bp repeat, with the (conserved) tRNA-Trp gene, followed by sequence currently unique to *N. brasiliensis*. Similarly, the shortest of the 5 corresponded to 90 copies of the tRNA-Ser gene interspersed with a *N. brasiliensis*-specific 80 bp spacer. In *C. elegans*, there are more than 600 tRNA genes. Some are clustered in small groups but never with such a massively repeated organization. The other 3 VeCTRs contain sequence that is not conserved. Remarkably, VeCTRs have recently been found in the genome of *C. elegans*, where they account for more than 1 Mb of sequence omitted from the current reference genome (E. Schwarz, personal communication) that was assembled primarily by Sanger sequencing of inserts from cosmid libraries [19]. Since they are not amenable to either short-read NGS or traditional cloning, they may have been overlooked in other species and potentially have a specific but as yet unknown biological role.

Returning to the analysis of the non-trivial sub-graphs, among the less complex ones, 28 linked just 3 contigs. Of these, 2 were due to the presence of VeCTRs and in 2, one contig matched the extremities of 2 non-overlapping contigs, but in a redundant manner (“Plain Link”) (**Figure 4A, B**). The others showed structures compatible with separate haplotypes. Of these, 12 had 2 contigs with homologous end sequences converging on a common contig (“contained”), a configuration that can also arise when base-calling errors are high. The remaining 12 had stronger support for being genuine haplotypes, including 3 sub-graphs with a bubble structure indicative of haplotype resolution, as supported by examination of the underlying reads (**Figure 4**). While attempting to identify a primary genomic haplotype might have some use in determining the true contiguity of the assembly, this assembly represents a community sample and preserves the observed read variation.

**Figure 4.**
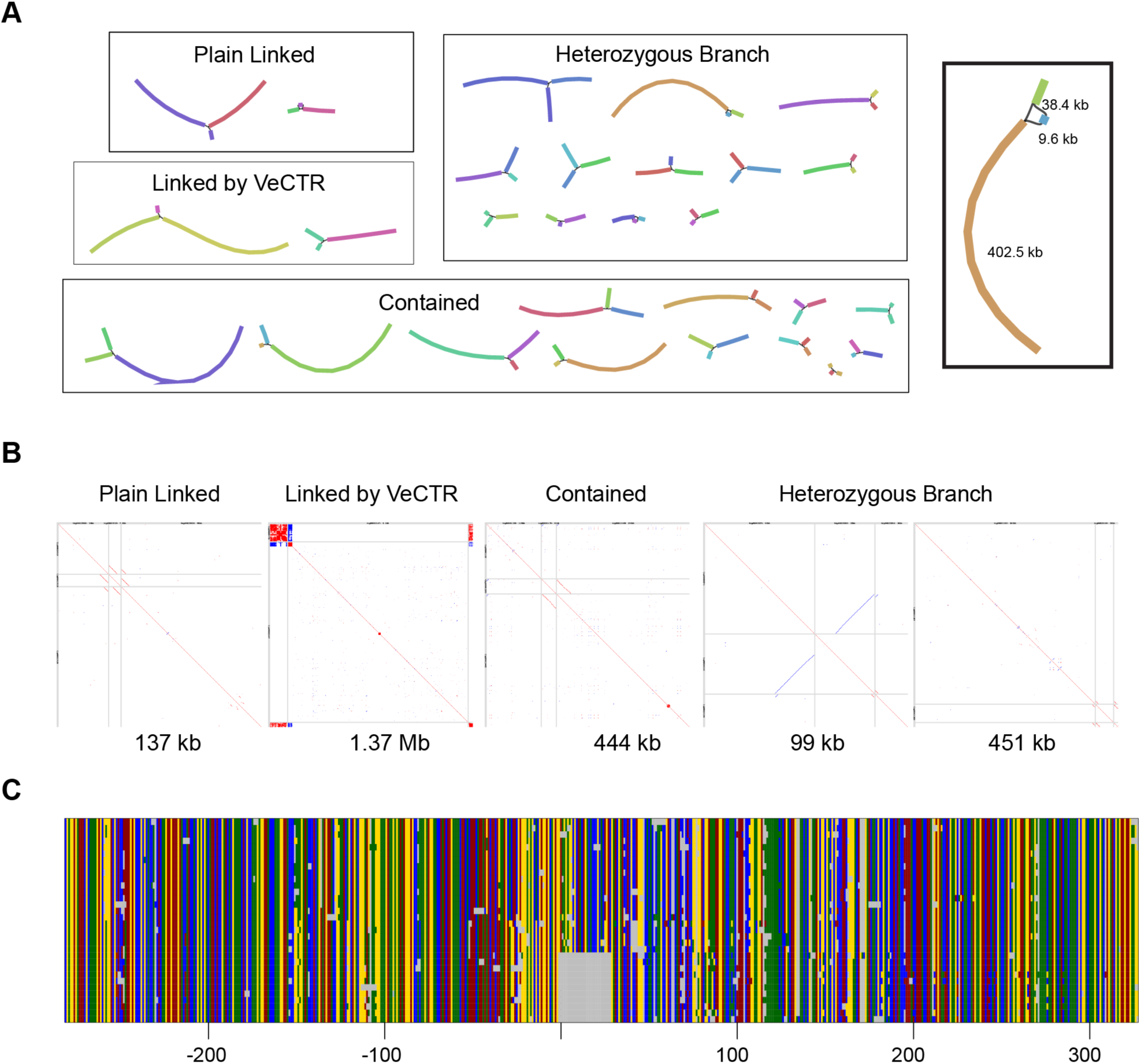
Classification of sub‐graphs and identification of a haplotype signature. (A)Bandage plots for the 28 simple GFA sub‐graphs made of 3 contigs. The box on the right is an enlarged view of a “heterozygous branch” sub‐graph with a total length of 451 kb. **(B)** Dot plots of LAST all‐against‐all minimum‐distance sequence comparisons between the 3 constituent contigs, delineated by the gray lines, for representatives for each of the sub‐graph classes. The sum of the contig lengths is indicated. The 451 kb sub‐graph is that enlarged in (A). **(C)** Multiple alignment of contig sequences from this sub‐graph and corresponding region of the DNA reads that map to them, 300 bp either side of the defining deletion. The color code is as in Figure 1A.

In addition to this capacity to uncover haplotypes within a population, overall, there was an impressive increase in the quality of the genome assembly compared to the current WTSI reference genome by standard contiguity criteria (**Figure 1F**, **Table 1**). The WTSI genome scored substantially higher (**Table 2**) when evaluated by Benchmarking Universal Single-Copy Orthologs (BUSCO) [20]. On the other hand, BLAST searches for the “missing” USCO genes identified plausible orthologues in most cases (results not shown). This suggested that even if the assembly continuity was improved, the local sequence quality was not sufficiently high to allow correct gene prediction by BUSCO. We therefore made use of a set of RNA-seq reads generated from our *N. brasiliensis* strain (GLG *et al*., unpublished) to correct the sequence of protein-coding genes, while not touching the genome assembly. We tried several methods, and settled on an approach based on genome-guided Trinity that gave the best results as judged by the markedly improved BUSCO scores (with 88% complete; **Table 3**). Indeed, following this correction the genome scored substantially higher than the WTSI reference, and close to the score of 91% complete BUSCO genes for the *de novo* assembly from the same RNA-seq reads. This indicates that the RNA-seq reads cover a substantial fraction of the transcriptome, and that the genome also has excellent coverage of most of the expressed *N. brasiliensis* genes. There was nonetheless an elevated proportion of fragmented USCOs (see below). We also noted a high proportion of duplicated USCOs. Inspection revealed that some of these were *bone fide* lineage-specific expansions. For example, the analysis uncovered 3 distinct loci encoding isoforms of fructose 1,6-bisphosphatase (PFAM: PF00316), as predicted also from the WTSI assembly. Pairs of USCOs were also found on homologous contigs. There were 4 such examples in the 12 “heterozygous branch” sub-graphs alone; this presumably reflects haplotypic variants.

**Table 3.** BUSCO scores following different methods of genome sequence correction. The percentage of complete (Comp) USCOs together with the number of single (Sin), duplicated (Dup), fragmented (Frag) or missing (Miss) USCO predicted from the uncorrected genome sequence, or the sequence after processing by different methods (see Methods for details). The figures for the WTSI reference are shown for comparison.

|  | Short BUSCO |  |  |  |  | Long BUSCO |  |  |  |  |
| --- | --- | --- | --- | --- | --- | --- | --- | --- | --- | --- |
| Description | Comp (%) | Sin (n) | Dup (n) | Frag (n) | Miss (n) | Comp (%) | Sin (n) | Dup (n) | Frag (n) | Miss (n) |
| Uncorrected | 49.2 | 437 | 46 | 128 | 371 | 65.0 | 585 | 53 | 122 | 222 |
| Bowtie2 + Pilon | 75.6 | 630 | 112 | 95 | 145 | 85.1 | 709 | 127 | 72 | 74 |
| HISAT2 + Pilon | 74.7 | 618 | 116 | 101 | 147 | 81.5 | 679 | 122 | 83 | 98 |
| Trinity<sup>#</sup><br>[*de novo*] | 90.8<br>90.7 | 681<br>441 | 211<br>450 | 65<br>64 | 25<br>27 | 91.3 | 446 | 451 | 59 | 26 |
| Trinity<sup>#</sup><br>[guided/Bowtie2] | 88.4<br>88.2 | 316<br>304 | 552<br>562 | 63<br>61 | 51<br>55 | 88.5 | 305 | 564 | 58 | 55 |
| Trinity<sup>#</sup><br>[filtered] | 87.9<br>87.7 | 644<br>613 | 219<br>248 | 58<br>60 | 61<br>61 | 88.3 | 618 | 249 | 55 | 60 |
| Trinity<sup>#</sup><br>[collapsed] | 87.2<br>87 | 786<br>791 | 71<br>64 | 58<br>63 | 67<br>64 | 87.6 | 796 | 64 | 57 | 65 |
| WTSI reference | 73.1 | 677 | 41 | 133 | 131 | 80.6 | 748 | 43 | 125 | 66 |

<sup>#</sup>Statistics in italics are from transcript mode BUSCO

While generating a complete high-quality annotation was beyond the scope of this study, we made use of expert knowledge regarding Carbohydrate-Active Enzymes (CAZymes) to provide a complementary insight into the predicted genes compared to the WTSI set [21]. Of the 62 different domain architectures among the 158 well-predicted CAZYme proteins from our assembly, only 36 were represented among the set of 96 such proteins in the WTSI reference proteome. Our assembly included a further 16 domain architectures represented among proteins that were flagged as being incorrectly predicted (N- or C-terminal fragments; (**Table S1**). Manual inspection revealed that in most cases, this was a consequence of the presence of non-tandem repeats (of which predicted transposable elements were a subset), where the repetition created ambiguity for transcript assembly (**Figure 5**). These structures, that also contributed to the presence of fragmented USCOs mentioned above, were even more of a problem for gene prediction in the WTSI assembly; they not only contributed to the fragmentation of USCOs and CAZymes (**Table S1**), but also introduced contig breaks that could not be bridged in the absence of long reads.

**Figure 5.**
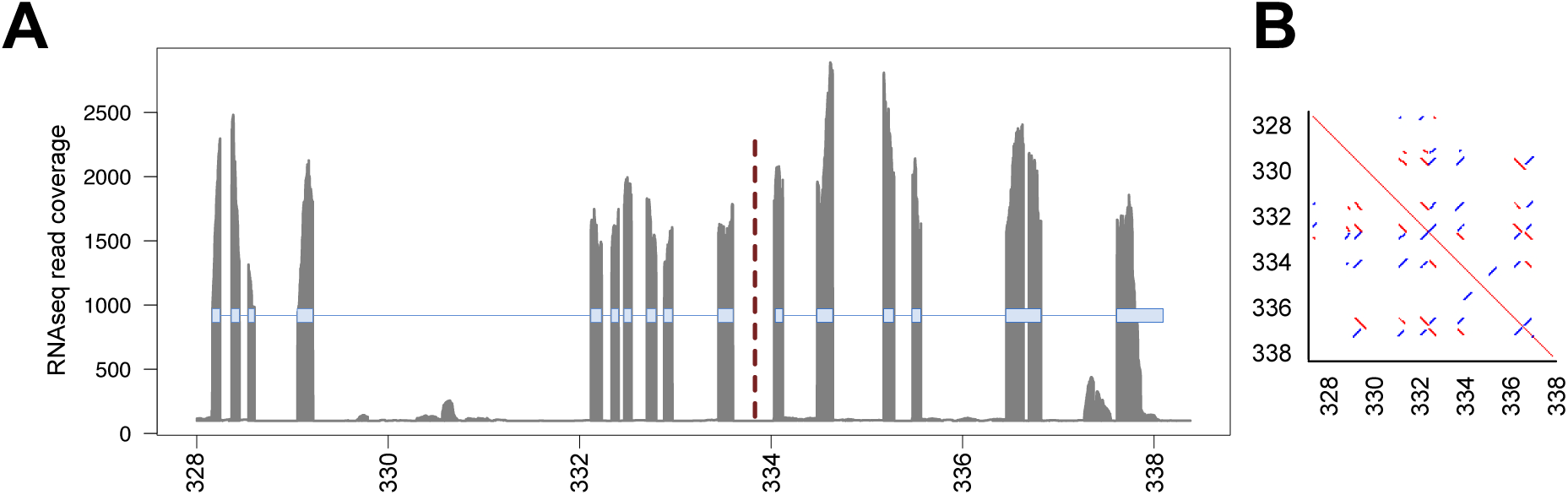
Repeat sequence confounds Trinity gene prediction. **(A)** CAZy analysis indicated that two adjacent Trinity‐generated transcripts (c16_g1_i2 and c16_g1_i1, left and right, respectively; exons indicated by blue boxes) had been incorrectly predicted since they correspond to C‐ and N‐terminal fragments of a single CAZyme gene (of the GT33 family). The read coverage of the exons for the two predicted transcripts was similar, supporting a single gene model. BUSCO erroneously called c16_g1_i1 as “complete” (EOG091H03EM0). **(B)** A self‐map dot‐plot of the genomic context in which these transcripts reside demonstrated a high degree of sequence similarity throughout the region, where the two predicted transcripts were split by a palindromic repeat motif that switched directions at the intersection of the two transcripts. The coordinates (in kbp) on the contig tig000206 are shown.

With a draft genome in hand, we then returned to the assembly we had generated from a single worm. Whole-genome amplification methods can suffer from chimera formation and amplification bias. The proportion of WGA reads that mapped over 90% of their length to the WGA assembly was lower than that of WGA reads mapping to the final assembly (41.9% vs 53.3%; **Table 4**), with a similar distribution of mapped read lengths, indicating that chimerism was not a significant problem here. Since the WGA reads that only mapped to the final assembly (in regions of low sequence coverage) represented 11% of the total number, sequencing depth was not a major limiting factor. As the stringent pairwise mapping also indicated that the WGA assembly captured a third (32.4%) of the predicted genome, it appears that genome amplification was indeed biased; future attempts should use alternate amplification (e.g. primer-free) approaches.

**Table 4.** Read mapping statistics. The number of reads that were generated from the whole genome amplified (WGA) DNA and that mapped to either the assembly made from those reads (WGA), or from those generated from unamplified DNA (Final) are shown in the left 2 columns. The number of reads that were generated from unamplified DNA (Unamplified) DNA and that mapped to either the assembly made from the WGA reads (WGA), or from those generated from unamplified DNA (Final) are shown in the right 2 columns. Since a similar fraction of WGA and unamplified reads do not map to the final assembly (ca. 10%), it appears that contaminanting DNA was effectively removed (see Methods for details).

| Read category | WGA |  | Unamplified |  |
| --- | --- | --- | --- | --- |
| Assembly | WGA | Final | WGA | Final |
| Read count | 1362775 | 1362775 | 2137475 | 2137475 |
| Mapped | 1106671 | 1233514 | 1525846 | 1922031 |
| Well-mapped<sup>1</sup> | 571180 | 727433 | 563302 | 1386713 |
| Uniquely-mapped<sup>2</sup> | 16977 | 143820 | 16568 | 412753 |
| Median mapped read proportion<sup>3</sup> | 91.1% | 94.1% | 65.6%<sup>4</sup> | 96.7% |
| Median mapped proportion excluding well-mapped reads | 56.1% | 65.9% | 31.1% | 67.2% |

<sup>1</sup> Reads that map at least 90% of their length to the reference assembly
<sup>2</sup> Reads that map only to the target assembly, and not the other assembly
<sup>3</sup> Median proportion of individual reads that aligned to the reference assembly
<sup>4</sup> Low mapping fraction partly due to the reduced haplotypic diversity in the WGA assembly

Together, our results support the conclusion that a *de novo* assembly of a high quality can be obtained using only long reads, even from a heterogeneous population, using a very modest sequencing depth (24X after trimming and correction). They also reveal the need for further improvements in resolving ambiguous contig architectures and in transcript-directed gene structure prediction. Nevertheless, since haplotypic variation could be detected even without RNA-seq-directed sequence correction, they clearly show the potential for using this approach to profile parasite populations, opening the way for detailed molecular epidemiological studies.

## Methods

### Canu 1.5

Changes to Canu relative to v1.4 are documented in the v1.5 release notes (available at https://github.com/marbl/canu/releases/tag/v1.5). Briefly, one major change was a switch in the alignment algorithm used to generate consensus for the corrected reads and final contigs. This improved the corrected reads by slightly increasing their identity and splitting fewer reads. Canu also started using the raw read overlaps to estimate corrected read lengths and used that information to inform its selection of reads for correction. As with all such analysis packages, bugs continue to be identified and corrected. The first release of v1.5, for example, included a bug that erroneously removed heterozygous edges such that in the type of sub-graphs shown in Figure 4A, the results presented would potentially correspond to lower bounds of complexity. This and other issues have been corrected in recent development versions of Canu and the latest public release.

### Nematode cultures

The *N. brasiliensis* strain used in this study was originally sourced from Lindsey Dent (University of Adelaide) and maintained at the Malaghan Institute for 22 years by serial passage through female Lewis rats [22]. Ethics approval is overseen and approved by the Victoria University of Wellington Animal Ethics Committee. The strain used for the Wellcome trust/Sanger Institute reference genome is not fully documented. In principal it derives from a line that had been maintained serially at NIMR Mill Hill by Bridget Ogilvie, starting in the early 1970s and then by Rick Maizel, at Imperial College. Murray Selkirk continued to maintain the strain at Imperial after R. Maizel moved to the University of Edinburgh, where he established parallel lines. These Edinburgh lines were supplemented on a number of occasions with cultures from Imperial College. The standard cycle involved injection of 4000-6000 L3 larvae into Sprague-Dawley rats, but was otherwise similar to that used at the Malaghan (R. Maizels, personal communication).

### DNA Preparation

Five samples of 25 mg each of frozen worms, cultured, harvested and purified as previously described [22] were sent to Simon Mayes (Oxford Nanopore Technologies) and subject to different methods of sample preparation (see **Table S2**).

#### 1. Sonication, DNeasy

Worms were disrupted by sonication and then processed using a Qiagen DNeasy kit (ATL / AL, 30min at 56°C). This yielded 900 ng DNA, 1800 ng RNA.

#### 2. Sonication, G2, Tip20

Worms were disrupted by sonication, then lysed using Qiagen buffer G2 (30 min at 56°C). The lysate was purified using a Qiatip 20 anion exchange column. This yielded 800 ng DNA.

#### 3. Direct lysis, DNeasy

Intact worms were added without prior disruption directly into Qiagen buffer G2 (30 mins at 56°C). The lysate was processed using a Qiagen DNeasy kit (ATL / AL, 30 min at 56°C). This yielded 700 ng DNA, 17700 ng RNA.

#### 4. Pipette squashed, G2, Tip20

Worms were mashed against the side of the sample tube using a pipette tip. The cells then underwent chemical lysis in the Qiagen buffer G2 (120 min at 56°C) and lysate was purified using the Qiatip 20 anion exchange column. The DNA was fragmented using a Covaris G-tube prior to library preparation. This yielded 1100ng DNA.

All samples were subsequently processed using the ONT 1D Ligation Sequencing Kit (SQK-LSK108), ligating the ONT adapter mix onto end-prepped and dA-tailed DNA. Each prepared library was loaded onto a different MinION flow cell for sequencing.

### Whole-Genome Amplification and Assembly Experimentation

DNA extracted from a single adult worm was amplified using a Qiagen Midi RepliG kit. Raw reads and called FASTQ files were obtained from ONT (called by workflow “1D RNN Basecalling 450bps FastQ v1.121”) following sequencing. Reads were filtered to exclude those from contaminating DNA using OneCodex [http://onecodex.com]. Reads that mapped to any DNA in the OneCodex database were excluded from the read set:

~~~
(zcat 0neCodex_RefSeq_132394.fastq.gz.results.tsv.gz | awk ‘{if($3 == 0){print $1}}’; zcat 0neCodex_0CD_132394.fastq.gz.results.tsv.gz | awk ‘{if($3 == 0){print $1}}’) | sort | uniq ‐d | gzip > 0Cunmapped_names_132394.txt.gz
pv 132394.fastq.gz | zcat | ∼/scripts/fastx-fetch.pl ‐i 0Cunmapped_names_132394.txt.gz | ∼/scripts/fastx-fetch.pl ‐v ‐i 0NTmapped_names_132394.txt.gz | gzip > 0Cunmapped_0NTunmapped_132394.fastq.gz
~~~

Filtered reads from the WGA sample were assembled using Canu v1.4, assuming a genome size of 150 Mb:

~~~
∼/install/canu/canu-1.4/Linux-amd64/bin/canu -nanopore-raw 0Cunmapped_0NTunmapped_132394.fastq.gz ‐p NbL5_0NTA ‐d NbL5_0NTA_testl genomeSize=150M
~~~

This is the WGA assembly presented in Figure 1B.

The WGA reads were then re-called using the updated basecaller (Albacore 1.1.0) that implements a time-domain correction (transducer) for homopolymeric regions in the called sequence, also available in the open-source Scrappie (https://github.com/nanoporetech/scrappie) basecaller:

~~~
read_fast5_basecaller.py ‐o fastq ‐i A_132394 ‐t 10 ‐s called_A_132394 ‐c r94_450bps_linear.cfg
~~~

Reads with a length of greater than 10k were extracted for subsequent analysis:

~~~
pv called_A_132394_albacore_1.1.0.fq.gz | zcat | ∼/scripts/fastx-fetch.pl ‐‐min 10000 | gzip > 10k_called_A_132394_albacore_1.1.0.fq.gz
~~~

In order to define the region of raw nanopore sequences for adapter exclusion, the >10k reads were mapped to 50M reads that had been generated by the Wellcome Trust Sanger Institute and that had been used for the existing WTSI assembly:

~~~
bowtie2 ‐p 10 ‐‐no‐unal ‐‐no‐mixed ‐‐local ‐x
10k_called_A_132394_albacore_1.1.0.fa ‐1 <(pv ∼/bioinf/MII^v^IR‐2017‐Dan‐01‐GBIS/GLG/ONT/aws/Sampled_50l^v^l_ERR063640.Rl.fq.gz | zcat) ‐2 ‘‐/bioinf/MIMR‐2017‐Dan‐01‐GBIS/GLG/ONT/aws/Sampled_50l^v^l_ERR063640.R2.fq.gz | samtools sort > WTSI_Sampled_50M_vs_10k_called_A_132394.bam
~~~

~~~
ææ find position of first mapped Illumina read for each nanopore read pv WTSI_Sampled_50M_vs_10k_called_A_132394.bam | samtools view ‐ | awk ‘{print $3j$4}’ | sort ‐k 1,1 ‐k 2,2n | sort ‐u ‐k 1,1 | gzip > firstHit_WTSI_Sampled_50M_vs_10k_called_A_132394.txt.gz
~~~

~~~
**##** determine position of last mapped Illumina read
pv WTSI_Sampled_50M_vs_10k_called_A_132394.bam | samtools view ‐ | awk
‘{print $3j$4}’ | sort ‐k 1,1 ‐k *2,2rn* | sort ‐u ‐k 1,1 | gzip >
lastHit_WTSI_Sampled_50M_vs_10k_called_A_132394.txt.gz
**##** count positions of first reads
zcat firstHit_WTSI_Sampled_50l^v^l_vs_10k_called_A_132394.txt.gz | awk ‘{print
$2}’ | sort ‐n | uniq ‐c | gzip >
firstBase_counts_WTSI_Sampled_50M_vs_10k_called_A_132394.txt.gz
~~~

When the 5’ end of Illumina reads mapped towards the start of nanopore reads, there was a common register shift of 28‐32 bases, corresponding to the presence of adapter sequences in the nanopore reads. To remove adapter sequences, a conservative 5’ trim of 65 bases was applied to both ends of all reads:

~~~
pv called_A_132394_albacore_1.1.0.fq.gz | zcat | ∼/scripts/fastx‐fetch.pl ‐‐min 1130 ‐‐max 1000000 | \
∼/scripts/fastx‐fetch.pl ‐t 65 | gzip > 65bpTrim_called_A_132394_albacore_l.1.0.fq.gz
~~~

Canu v1.5 was used to assemble the trimmed reads. The assembly was done in stages (with an assembly at each stage) to determine whether or not particular stages were redundant for the assembly:

~~~
**##** attempt assembly‐only with Canu v1.5
∼/install/canu/canu‐1.5/Linux‐amd64/bin/canu ‐assemble ‐nanopore‐raw
65bpTrim_called_A_132394_albacore_1.1.0.fq.gz ‐p Nb_ONTA_65bpTrim_tl ‐d
Nb_ONTA_65bpTrim_tl genomeSize=300M
**##** attempt assembly + correction
∼/install/canu/canu‐1.5/Linux‐amd64/bin/canu ‐assemble ‐nanopore‐corrected
65bpTrim_called_A_132394_albacore_1.1.0.fq.gz ‐p Nb_0NTA_65bpTrim_t2 ‐d
Nb_0NTA_65bpTrim_t2 ‐correct genomeSize=300M
**##** attempt stringent trim with corrected reads
∼/install/canu/canu‐1.5/Linux‐amd64/bin/canu ‐trim‐assemble ‐p
Nb_ONTA_65bpTrim_t3 ‐d Nb_ONTA_65bpTrim_t3 genomeSize=300M ‐nanopore‐
corrected ext-link-type="uri" xlink:href="http://Nb_0NTA_65bpTrim_t2/Nb_0NTA_65bpTrim_t2.correctedReads.fasta.gz">Nb_0NTA_65bpTrim_t2/Nb_0NTA_65bpTrim_t2.correctedReads.fasta.gz ‐trim‐assemble trimReadsOverlap=500 trimReadsCoverage=5 obtErrorRate=0.25
~~~

An alternative, less‐stringent overlap was also attempted (with trimReadsCoverage=2), but resulted in a less complete assembly. The results of analysis surrounding the different assemblies suggested that the default Canu assembly process of correction, trimming, then assembly produced the best outcome, namely the WGA assembly presented in Figure 1C.

### Genome Assembly

Raw reads and called FASTQ files were obtained from ONT (called by workflow “1D RNN Basecalling 450bps FastQ v1.121”) following sequencing. The raw reads were re‐called using Albacore 1.1.0:

~~~
for x in $(ls ‐d [CFED]_??????); do echo ${x}; read_fast5_basecaller.py ‐t 6 ‐i ${x} ‐s called_${x} ‐o fastq ‐c r94_450bps_linear.cfg; done
~~~

Reads were trimmed by 65bp at each end to exclude adapters, then Canu v1.5 was run with default parameters, assuming a genome size of 300 Mb:

~~~
**##** trim reads
pv called_[CFED]_*_albacore_1.1.0.fq.gz | zcat | ∼/scripts/fastx‐fetch.pl ‐t
65 | gzip > called_CFED_65bptrim_albacore_1.1.0.fq.gz
## run Canu
∼/install/canu/canu‐1.5/Linux‐amd64/bin/canu ‐nanopore‐raw
called_CFED_65bptrim_albacore_1.1.0.fq.gz ‐p Nb_ONTCFED_65bpTrim_tl ‐d
Nb_ONTCFED_65bpTrim_tl genomeSize=300M
~~~

Bowtie2 was used in local mode to map RNA‐seq reads to the assembled genome contigs:

~~~
bowtie2 ‐p 10 ‐‐local ‐x Nb_ONTCFED_65bpTrim_tl.contigs.fasta ‐1 ../1563‐all_Rl_trimmed.fastq.gz ‐2 ../1563‐all_R2_trimmed.fastq.gz | samtools sort > 1563_vs_uncorrected_NOCFED.bam
~~~

Pilon was used to correct based on the RNA‐seq mapping to the genome, with structural reassembly disabled (in case it collapsed introns):

~~~
Java ‐Xmx40G ‐jar ∼/install/pilon/pilon‐1.22.jar ‐‐genome
Nb_ONTCFED_65bpTrim_tl.contigs.fasta ‐‐frags 1563_vs_uncorrected_NOCFED.bam ‐‐fix snpSjindels ‐‐output BT2Pilon_N0CFED ‐‐gapmargin 1 ‐‐mingap 10000000 ‐‐threads 10 ‐‐changes 2 &amp;gt;BT2Pilon_N0CFED.stderr.txt l &amp;gt;BT2Pilon_N0CFED.stdout.txt
~~~

Contigs that were entirely composed of homopolymer sequences were identified using grep and removed from the assembly:

~~~
**##** identify homopolymer (and binary division‐rich) regions pv BT2Pilon_N0CFED.fasta | ∼/scripts/fastx‐hplength.pi > hplength_BT2Pilon_N0CFED.txt
pv BT2Pilon_N0CFED.fasta | ∼/scripts/fastx‐hplength.pi ‐mode YR > hplength_YR_BT2Pilon_N0CFED.txt
pv BT2Pilon_N0CFED.fasta | ∼/scripts/fastx‐hplength.pi ‐mode SW > hplength_SW_BT2Pilon_N0CFED.txt
pv BT2Pilon_N0CFED.fasta | ∼/scripts/fastx‐hplength.pi ‐mode MK > hplength_MK_BT2Pilon_N0CFED.txt ## example grep hunt for repeated sequence
cat BT2Pilon_N0CFED.fasta | grep ‐e ‘ˆ[AT]\{80\}’ ‐e ‘ˆ >‘ | grep ‐‐no‐group‐separator ‐B 1 ‘ˆ[AT]\{80\}’ | ∼/scripts/fastx‐length.pl ## exclude contigs and sort by length
∼/scripts/fastx‐fetch.pl ‐v tig00010453 tig00024413 tig00024414 tig00023947 | ∼/scripts/fastx‐sort.pi ‐1 > Nb_ONTCFED_65bpTrim_tl.contigs.hpcleaned.fasta
~~~

This produced the final assembly described in the paper. At this stage, we had an assembled genome, but validation of the genome was difficult. We decided to carry out a draft genome‐guided transcriptome assembly with Trinity. To evalaute the completeness of the genome, we focused on expressed genes and used a set of Illumina RNA‐seq reads to perform a genome‐guided transcriptome assembly using Trinity.

The RNA‐seq reads were remapped to the corrected assembly for genome‐guided Trinity:

~~~
bowtie2 ‐p 10 ‐t ‐‐local ‐‐score‐min 6,20,8 ‐p 10 ‐x BT2Pilon_N0CFED_hpcleaned.fasta ‐‐rf ‐X 15000 ‐1 \
../1563‐all_Rl_trimmed.fastq.gz ‐2 <(pv ../1563‐all_R2_trimmed.fastq.gz | zcat) 2 &amp;gt;bowtie2_1563_vs_BNOCFED_hp.summary.txt | samtools sort > bowtie2_1563_vs_BNOCFED_hp.bam
**##** Trinity assembly; assume introns can be up to 15kb in length ∼/install/trinity/trinityrnaseq‐Trinity‐v2.4.0/Trinity ‐‐CPU 10 ‐‐genome_guided_bam bowtie2_1563_vs_BNOCFED_hp.bam ‐‐genome_guided_max_intron 15000 ‐‐max_memory 40G ‐‐SS_lib_type RF ‐‐output trinity_BNOCFED
~~~

The assembly that Trinity generated had similar completeness (as measured by BUSCO) to a *de novo* assembly generated using the same RNA‐seq reads (see **Table 3**). The assembly was, however, very large, with over 350k contigs (see **Table 1**), most likely due to isoform fragments being included in the transcriptome. We carried out additional filtering steps, reducing the number of assembled contigs while maintaining similar BUSCO completeness scores.

### Transcript Filtering

The estimated numbers of mapped RNA‐seq reads (as predicted by Salmon [DOI 10.1038/nmeth.4197]) were used to filter transcripts, because the genome‐guided assembly was based on RNA‐seq expression. RNA‐seq reads were mapped to the Bowtie2/Pilon genome‐guided assembly, and a threshold of 50 reads for true expression of complete BUSCO genes was chosen for filtering transcripts from the genome‐guided Trinity transcriptome. The longest ORF from each transcript was extracted to generate a protein sequence for further collapsing using cd‐hit to remove protein sequences that had at least 98% identity to a longer protein, leaving 56980 transcripts. Each of these steps are outlined below.

The RNA‐seq reads were mapped to the Trinity‐generated transcripts using Salmon:

~~~
**##** create Salmon index
∼/install/salmon/Salmon‐0.8.2_linux_x86_64/bin/salmon index ‐t Trinity‐BNOCFED.fasta ‐i Trinity‐BNOCFED.fasta.sai
**##** quantify transcript coverage with Salmon
∼/install/salmon/Salmon‐0.8.2_linux_x86_64/bin/salmon quant ‐i Trinity‐BNOCFED.fasta.sai ‐1 ../../1563‐all_Rl_trimmed.fastq.gz ‐2 ../../1563‐all_R2_trimmed.fastq.gz ‐p 10 ‐o quant/1563‐all_quant ‐1 A
~~~

The expression of BUSCO genes was used to set a credible signal cutoff (**Figure S3**). A threshold of 50 mapped reads was chosen, as 98% of complete BUSCO sequences had more than 50 mapped reads. The transcripts were subsetted based on their expression scores:

~~~
**##** Subset transcripts based on a threshold of 50 counts pv Trinity‐GG.fasta | ∼/scripts/fastx‐fetch.pl ‐i HighCount_Transcripts_Num50.txt > HighCount50_TBNOCFED.fasta
~~~

The longest Met to Stop ORF was identified for each transcript for the purpose of protein‐based clustering with CDHIT:

~~~
pv HighCount50_TBNOCFED.fasta | getorf ‐find 1 ‐noreverse ‐sequence
/dev/stdin ‐outseq /dev/stdout | ∼/scripts/fastx‐isofilter.pl ‐о >
longest_MetStopORF_HC50_TBNOCFED.fasta
**##** run cdhit
cdhit ‐T 10 ‐c 0.98 ‐i longest_MetStopORF_HC50_TBNOCFED.fasta ‐о
cdhit_0.98_LMOHC50_TBNOCFED.prot.fasta
**##** identify names of longest representative proteins for each cluster
grep ‘ˆ >‘ cdhit_0.98_LMOHC50_TBNOCFED.prot.fasta | perl ‐pe ‘s/ˆ >//’ >
cdhit_0.98_LMOHC50_TBNOCFED.prot.names.txt
**##** fetch transcripts associated with the representative proteins
pv HighCount50_TBNOCFED.fasta | ∼/scripts/fastx‐fetch.pl ‐i
cdhit_0.98_LMOHC50_TBNOCFED.prot.names.txt >
cdhit_0.98_LMOHC50_TBNOCFED.tran.fasta
~~~

The longest isoform for each gene was identified, producing an isoform‐collapsed transcriptome subset, which was clustered at the protein level by CDHIT at 90% identity:

~~~
**##** extract longest protein for each transcript
cat cdhit_0.98_LMOHC50_TBNOCFED.prot.fasta | ∼/scripts/fastx‐isofilter.pl >
LI_CD98LMOHC50_TBNOCFED.prot.fasta
**##** cluster at 90% via CDHIT
cdhit ‐i LI_CD98LMOHC50_TBNOCFED.prot.fasta ‐о
CDLI_CD98LMOHC50_TBNOCFED.prot.fasta
**##** find associated transcripts
grep ‘ˆ >‘ CDLI_CD98LMOHC50_TBNOCFED.prot.fasta | perl ‐pe ‘s/ˆ >//’ >
CDLI_CD98LMOHC50_TBNOCFED.names.txt
cat cdhit_0.98_LMOHC50_TBNOCFED.tran.fasta | ∼/scripts/fastx‐fetch.pl ‐i
CDLI_CD98LMOHC50_TBNOCFED.names.txt > CDLI_CD98LMOHC50_TBNOCFED.tran.fasta
~~~

BUSCO was run on the collapsed transcripts to provide one measure of genome completeness:

~~~
**##** run BUSCO on isoform‐collapsed transcripts in long genome mode python ∼/install/busco/BUSCO.py ‐i ../CDLI_CD98LMOHC50_TBNOCFED.tran.fasta ‐о BUSCO_longgeno_CDLI_CD98LMOHC50_TBNOCFED_nematodes ‐1 \ ∼/install/busco/nematoda_odb9 ‐m geno ‐c 10 ‐‐long
**##** run BUSCO on isoform‐collapsed transcripts in transcript mode python ∼/install/busco/BUSCO.py ‐i ../CDLI_CD98LMOHC50_TBNOCFED.tran.fasta ‐о BUSCO_tran_CDLI_CD98LMOHC50_TBNOCFED_nematodes ‐1 \ ∼/install/busco/nematoda_odb9 ‐m tran ‐c 10
~~~

This filtered set had very similar BUSCO scores to the original genome‐guided Trinity assembly, despite reducing the number of transcripts down to 1/6 of their original count, and the length of the transcriptome down to less than a third of its original size. The number of duplicated BUSCO genes in the filtered set suggests that this set could probably be made smaller with a looser cd‐hit‐est clustering, although there is a chance that such a reduction may cause gene copies to be clustered together.

## Acknowledgements

This study was supported by program grant funding from the Health Research Council of New Zealand (14/003) and the Marjorie Barclay Trust. SK was supported by the Intramural Research Program of the National Human Genome Research Institute, National Institutes of Health. We thank Rick Maizel for discussion, Simon Mayes and colleagues at Oxford Nanopore Technologies for their assistance. This study utilized the computational resources of the Biowulf system at the National Institutes of Health, Bethesda, MD (https://biowulf.nih.gov).

### Data Availability

MinION reads (base‐called FASTQ and raw) used for this assembly project (as well as RNASeq reads used for assembly correction) are available from ENA, project PRJNA328296.

Accessory scripts used for sequence and assembly analysis (see Methods) can be found at https://github.com/gringer/bioinfscripts.

---

**Figure S1.**
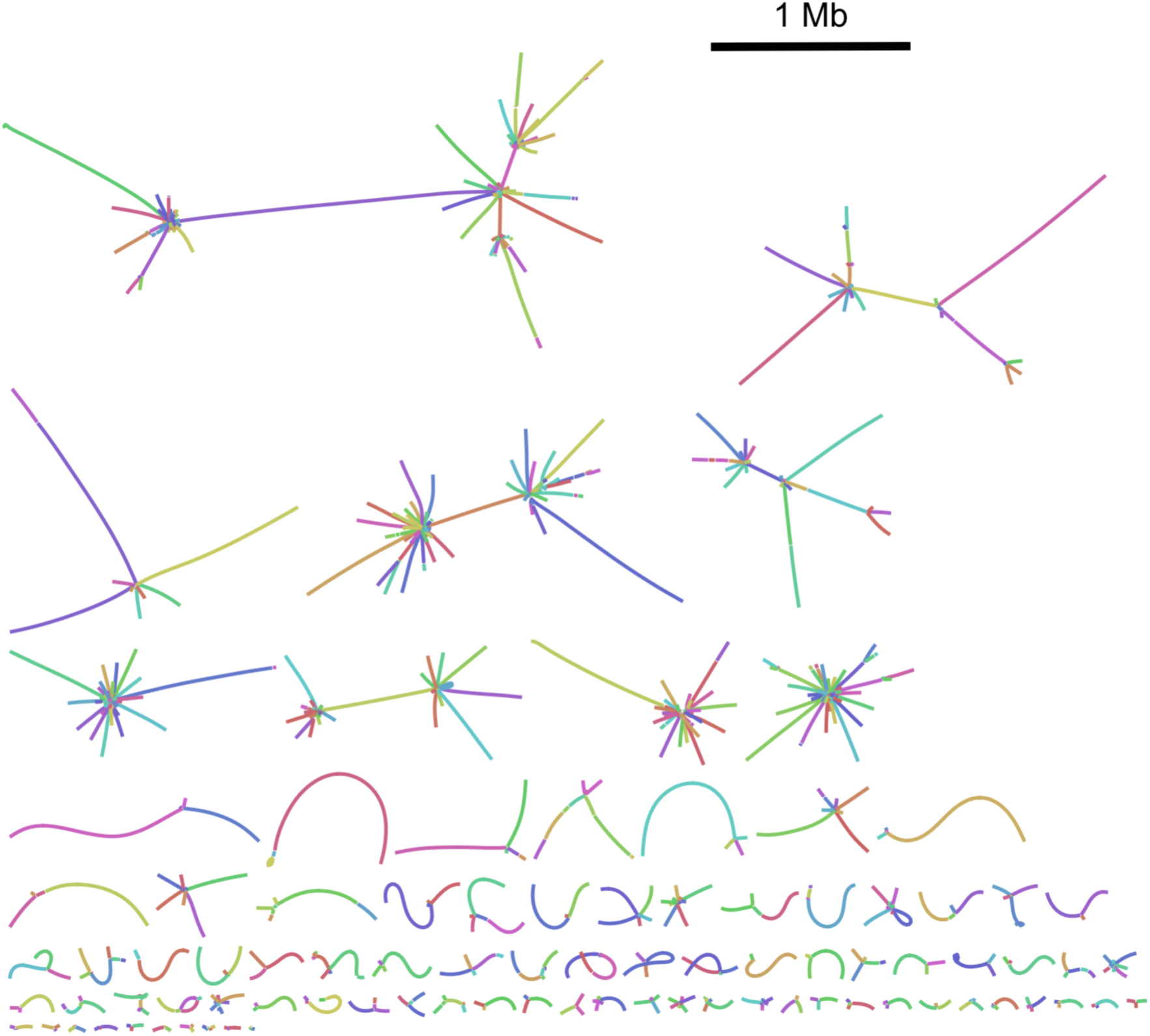
Branched graphs for the genome assembly from unamplified DNA. The largest one (top left) involved 106 contigs for a total of 13.3 Mb.

**Figure S2.**
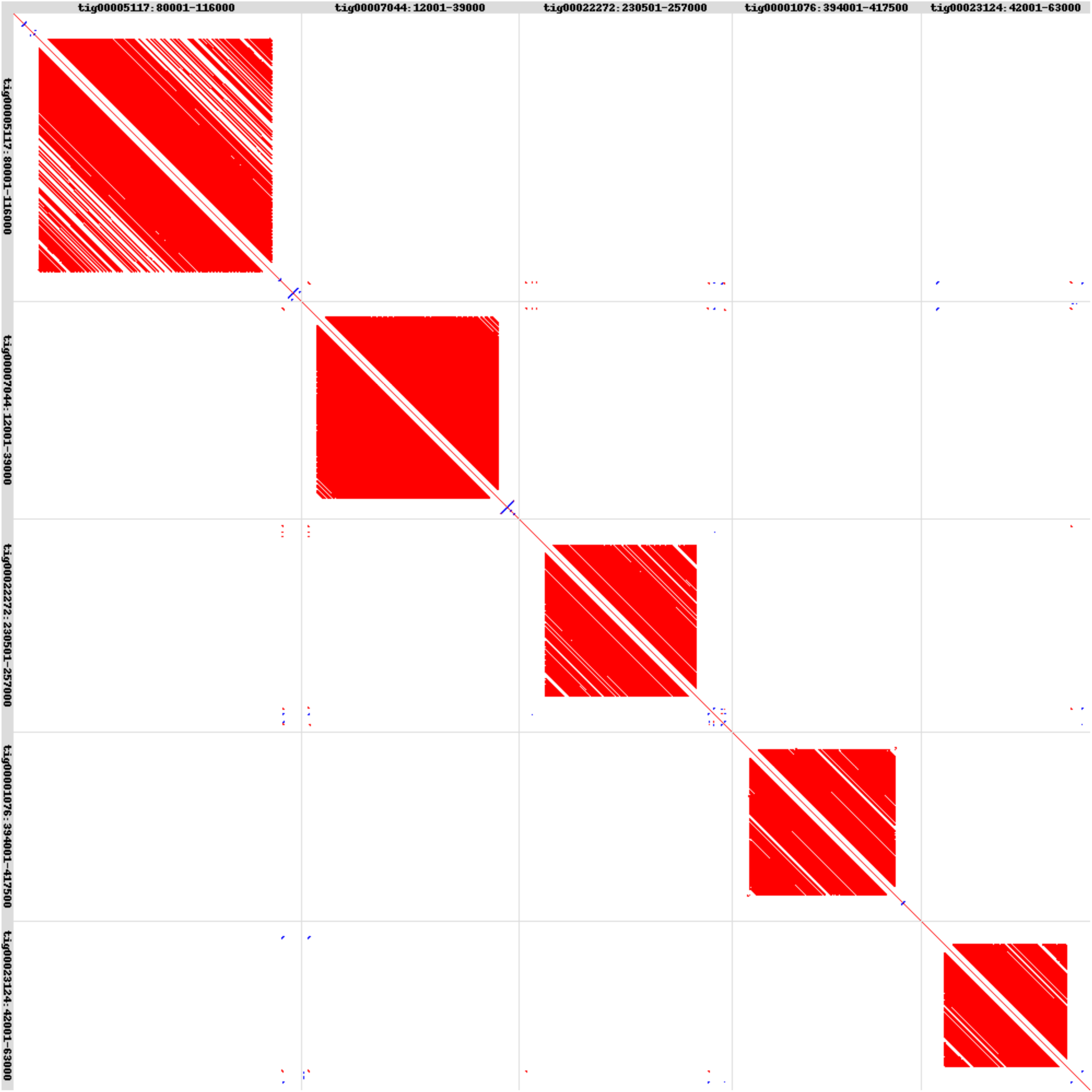
Dot plots of all‐against all sequence comparisons between the 5 most compressible VeCTR regions, based on minimum‐distance alignments using LAST‐align, created using LAST‐dotplot. The VeCTRs, with their unique flanking sequences are arranged in order of size (36 kb, 27 kb, 26.5 kb, 23.5 kb, 21 kb). The longest of these 5 VeCTRs corresponds to 147 repeats of tRNA‐Trp followed by 114 bp of non‐conserved sequence, while the shortest contains 90 copies of tRNA‐Ser with 80 bp of non‐conserved sequence.

**Figure S3.**
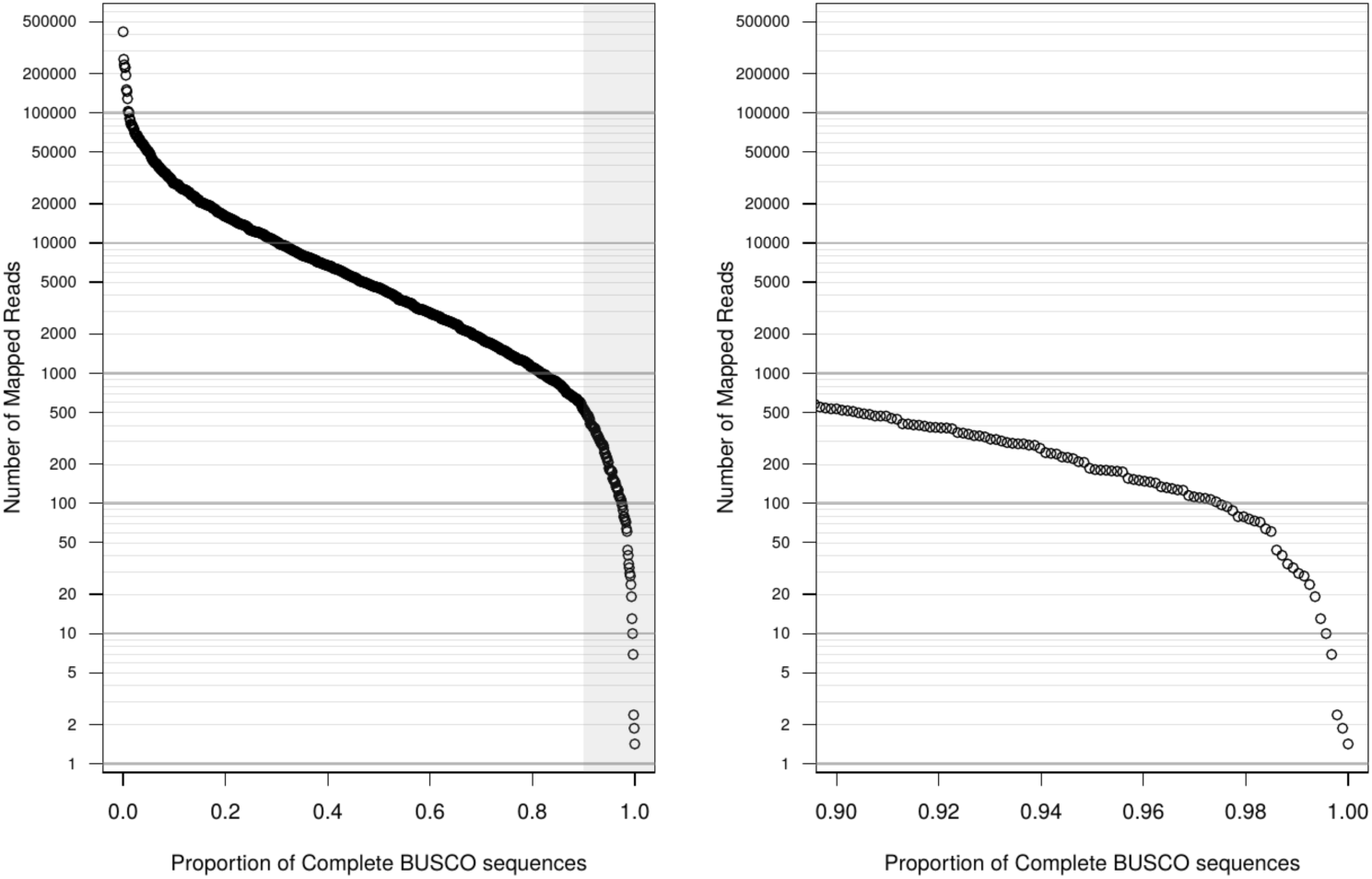
The distribution of the number of mapped reads for BUSCO sequences. The area in gray on the left‐hand graph is shown enlarged on the right.

## References

1. Wit J, Gilleard JS: Resequencing Helminth Genomes for Population and Genetic Studies. Trends Parasitol 2017, 33(5):388‐399.

2. Sotillo J, Sanchez‐Flores A, Cantacessi C, Harcus Y, Pickering D, Bouchery T, Camberis M, Tang SC, Giacomin P, Mulvenna J et al: Secreted proteomes of different developmental stages of the gastrointestinal nematode Nippostrongylus brasiliensis. Mol Cell Proteomics 2014, 13(10):2736‐2751.

3. Koren S, Schatz MC, Walenz BP, Martin J, Howard JT, Ganapathy G, Wang Z, Rasko DA, McCombie WR, Jarvis ED et al: Hybrid error correction and de novo assembly of single‐molecule sequencing reads. Nat Biotechnol 2012, 30(7):693‐700.

4. Zimin AV, Puiu D, Luo MC, Zhu T, Koren S, Marcais G, Yorke JA, Dvorak J, Salzberg SL: Hybrid assembly of the large and highly repetitive genome of *Aegilops tauschii*, a progenitor of bread wheat, with the MaSuRCA mega‐reads algorithm. Genome Res 2017, 27(5):787‐792.

5. Ye C, Hill CM, Wu S, Ruan J, Ma ZS: DBG2OLC: Efficient Assembly of Large Genomes Using Long Erroneous Reads of the Third Generation Sequencing Technologies. Sci Rep 2016, 6:31900.

6. Jansen HJ, Liem M, Jong‐Raadsen SA, Dufour S, Weltzien FA, Swinkels W, Koelewijn A, Palstra AP, Pelster B, Spaink HP et al: Rapid de novo assembly of the European eel genome from nanopore sequencing reads. Sci Rep 2017, 7(1):7213.

7. Berlin K, Koren S, Chin CS, Drake JP, Landolin JM, Phillippy AM: Assembling large genomes with single‐molecule sequencing and locality‐sensitive hashing. Nat Biotechnol 2015, 33(6):623‐630.

8. Li H: Minimap and miniasm: fast mapping and de novo assembly for noisy long sequences. Bioinformatics 2016, 32(14):2103‐2110.

9. Chin CS, Peluso P, Sedlazeck FJ, Nattestad M, Concepcion GT, Clum A, Dunn C, O’Malley R, Figueroa‐Balderas R, Morales‐Cruz A et al: Phased diploid genome assembly with single‐molecule real‐time sequencing. Nat Methods 2016, 13(12):1050‐1054.

10. Chin CS, Alexander DH, Marks P, Klammer AA, Drake J, Heiner C, Clum A, Copeland A, Huddleston J, Eichler EE et al: Nonhybrid, finished microbial genome assemblies from long‐read SMRT sequencing data. Nat Methods 2013, 10(6):563‐569.

11. Koren S, Harhay GP, Smith TP, Bono JL, Harhay DM, McVey SD, Radune D, Bergman NH, Phillippy AM: Reducing assembly complexity of microbial genomes with single‐molecule sequencing. Genome Biol 2013, 14(9):R101.

12. Koren S, Phillippy AM: One chromosome, one contig: complete microbial genomes from long‐read sequencing and assembly. Curr Opin Microbiol 2015, 23:110‐120.

13. Gordon D, Huddleston J, Chaisson MJ, Hill CM, Kronenberg ZN, Munson KM, Malig M, Raja A, Fiddes I, Hillier LW et al: Long‐read sequence assembly of the gorilla genome. Science 2016, 352(6281):aae0344.

14. Bickhart DM, Rosen BD, Koren S, Sayre BL, Hastie AR, Chan S, Lee J, Lam ET, Liachko I, Sullivan ST et al: Single‐molecule sequencing and chromatin conformation capture enable de novo reference assembly of the domestic goat genome. Nat Genet 2017, 49(4):643‐650.

15. Jarvis DE, Ho YS, Lightfoot DJ, Schmockel SM, Li B, Borm TJ, Ohyanagi H, Mineta K, Michell CT, Saber N et al: The genome of *Chenopodium quinoa*. Nature 2017, 542(7641):307‐312.

16. Koren S, Walenz BP, Berlin K, Miller JR, Bergman NH, Phillippy AM: Canu: scalable and accurate long‐read assembly via adaptive k‐mer weighting and repeat separation. Genome Res 2017, 27(5):722‐736.

17. Lu H, Giordano F, Ning Z: Oxford Nanopore MinION Sequencing and Genome Assembly. Genomics Proteomics Bioinformatics 2016, 14(5):265‐279.

18. Subirana JA, Messeguer X: A satellite explosion in the genome of holocentric nematodes. PLoS One 2013, 8(4):e62221.

19. The C. elegans sequencing consortium: Genome sequence of the nematode C. elegans: a platform for investigating biology. Science 1998, 282(5396):2012‐2018.

20. Simao FA, Waterhouse RM, Ioannidis P, Kriventseva EV, Zdobnov EM: BUSCO: assessing genome assembly and annotation completeness with single‐copy orthologs. Bioinformatics 2015, 31(19):3210‐3212.

21. Lombard V, Golaconda Ramulu H, Drula E, Coutinho PM, Henrissat B: The carbohydrate‐active enzymes database (CAZy) in 2013. Nucleic Acids Res 2014, 42(Database issue):D490‐495.

22. Chandler J, Camberis M, Bouchery T, Blaxter M, Le Gros G, Eccles DA: Annotated mitochondrial genome with Nanopore R9 signal for *Nippostrongylus brasiliensis*. F1000Res 2017, 6:56.

23. Wintersinger JA, Wasmuth JD: Kablammo: an interactive, web‐based BLAST results visualizer. Bioinformatics 2015, 31(8):1305‐1306.

